# Timing matters: exon skipping therapy is most effective when initiated early in a mouse model of Duchenne muscular dystrophy

**DOI:** 10.64898/2025.12.23.696144

**Authors:** Sofia Stenler, Junyu Huang, Tirsa L.E. van Westering, Anna M.L. Coenen-Stass, Kaarel Krjutškov, Samir EL Andaloussi, Matthew J.A. Wood, Thomas C. Roberts

## Abstract

Exon skipping is a leading therapeutic approach for Duchenne muscular dystrophy (DMD), a progressive muscle wasting disorder caused by pathogenic variants in the *DMD* gene that typically disrupt the translation reading frame. This approach aims to modulate *DMD* pre-mRNA splicing to re-frame the transcript and generate an internally deleted but partially functional quasi-dystrophin protein. Four exon skipping drugs have received FDA accelerated approval, although their clinical efficacy is very limited. To investigate how treatment timing influences exon skipping outcomes, dystrophin-deficient *mdx* mice were injected with peptide-phosphorodiamidate morpholino oligonucleotide (PPMO) exon skipping conjugates beginning at either the adult (12-week) or aged (75-week) stages, and tibialis anterior muscles harvested for biochemical and transcriptomic analyses. Mean *Dmd* exon 23 skipping was 79% in adults and 44% in aged PPMO-treated *mdx* mice, whereas dystrophin protein restoration was 35% and 8%, respectively. Histopathological improvements were only evident in the adult-treated mice. PPMO-treatment in adult *mdx* mice induced a broad transcriptomic shift towards a wild-type signature, whereas treatment in aged mice resulted in negligible gene expression changes, indicating that late intervention is ineffective at reversing disease-associated pathologies despite low-level dystrophin restoration. Dystrophin transcript imbalance was corrected only in adult-treated *mdx* mice. Increased expression of the dystrophin-repressing microRNA miR-31-5p, which was more strongly upregulated in aged *mdx* muscle, provides a potential mechanistic explanation. In conclusion, PPMO-mediated exon skipping is substantially more effective when initiated in adult rather than aged dystrophic muscle, supporting early therapeutic intervention in DMD-affected individuals.

## Introduction

Duchenne muscular dystrophy (DMD) is an X-linked, monogenic muscle wasting disorder caused by pathogenic variants in the gene encoding the dystrophin protein (human: *DMD*, mouse: *Dmd*) with an incidence of 1 in 5,000 live male births.^1^ Dystrophin is a structural and signalling protein which acts to assemble a complex of proteins at the sarcolemma (the dystrophin-associated protein complex, DAPC). Loss of dystrophin sensitizes muscle to contractile damage,^2^ leading to cycles of muscle turnover (i.e. myonecrosis and compensatory regeneration) and persistent inflammation. As a result, muscle quality declines over time, with progressive replacement of myofibers with connective and adipose tissue (i.e. fibro-fatty degeneration). DMD patients typically present with signs of muscle weakness, delays in the achievement of motor milestones, and movement difficulties. Loss of ambulation typically occurs around the age of ten-years, and DMD is ultimately fatal, as a consequence of cardiac or respiratory failure around age thirty.^3–5^ Standard-of-care treatment of DMD involves long-term multidisciplinary care^6–8^ and corticosteroid therapy, the latter of which has been demonstrated to delay the loss of ambulation by a few years^7,9,10^ but is associated a number of undesirable side effects^11^ As such, DMD remains a substantial health burden, underscoring the need for therapies that meaningfully modify disease progression.

Recent years have seen substantial progress in the field of experimental therapeutics for DMD.^12^ Various strategies have been employed to restore dystrophin. For example, antisense oligonucleotide (ASO)-mediated exon skipping can be utilised to modulate alternative splicing such that the dystrophin translation reading frame is restored.^13–15^ Four such antisense oligonucleotide drugs received accelerated approval from the US FDA for the treatment of DMD boys (Eteplirsen, Viltolarsen, Golodiresen, and Casimersen). While a major achievement for the DMD research and patient communities, the dystrophin restoration achieved by these compounds is minimal,^16–19^ and of debatable clinical benefit.^20–25^ Similarly, the FDA approved the first gene therapy for DMD (Elevidys) in 2023,^26^ despite failure to reach its clinical endpoint in a Phase 3 trial.^27^ Furthermore, the safety of Elevidys (and high dose adeno-associated virus-based gene therapies in general) has been called into question following multiple patient deaths.^28^

The current generation of exon skipping ASOs are all phosphorodiamidate morpholino oligonucleotides (PMOs). This chemistry does not occur in nature, and is widely regarded as being safe.^29^ However, PMO delivery to skeletal (and especially cardiac) muscle is very limited, which has motivated a search for delivery-assisting moieties that can enhance their activity. Our group, and others, has extensively explored the use of peptide-conjugated PMOs (PPMOs) for such purposes.^30–33^ PPMOs are much more potent than unconjugated PMOs, and exhibit exon skipping activity in cardiac tissue.^30–33^ PPMOs are currently being commercially developed by PepGen Ltd and Entrada Therapeutics.^34,35^ Similarly, Avidity Biosciences and Dyne Therapeutics are developing PMO-antibody and PMO-FAb fragment therapeutics for DMD.^36,37^

The *mdx* mouse is the most commonly-used murine model of DMD, which carries a premature termination codon in *Dmd* exon23, skipping of which results in an X-linked muscular dystrophy that exhibits some aspects of DMD pathology.^38,39^ We have previously performed a variety of omics analyses in this mouse with a focus on the effects of dystrophin restoration therapies, including gene expression/microRNA microarray,^40,41^ small RNA-Seq,^42^ and high resolution mass spectrometry-based proteomics.^40,43^

A number of factors are important when considering the efficacy of dystrophin restoration therapies. The total amount of dystrophin restored. It is widely accepted that the more dystrophin that can be restored the better. Various lines of evidence suggest that ∼10% of healthy dystrophin levels is a sensible threshold for therapeutic benefit.^44–46^ (ii) The quality of dystrophin restored. Exon skipping splice correction and microdystrophin gene therapies both rely on the generation of internally-deleted quasi-dystrophin proteins that resemble those found in some Becker muscular dystrophy patients,^47^ and which are associated with mild pathology. However, such quasi-dystrophins are unlikely to fully compensate for the loss of the full-length dystrophin protein. There is a growing appreciation that full-length dystrophins are desirable, which presents a challenge given that the size of the dystrophin ORF (∼11 kb) exceeds the packaging capacity of AAV vectors (∼4.6 kb). Restoration of full-length dystrophin has been demonstrated in human clinical trial participants treated with an exon skipping therapy to treat DMD exon2 duplication.^48^ Furthermore, there are multiple other efforts to develop full-length dystrophin delivery strategies using split-vector approaches,^49–51^ or by using vectors with higher packaging capacities.^52–54^ (iii) Correct localization of dystrophin at the sarcolemma. Work from our group, and others, has shown that uniform distribution of dystrophin protein along the sarcolemma within a myofiber is an important determinant of therapeutic success.^55–59^ Importantly, exon skipping treatment with PPMO conjugates induced a uniform pattern of sarcolemmal dystrophin coverage (even at low doses),^56^ while CRISPR-Cas9-mediated exon deletion resulted in a patchy pattern of dystrophin distribution.^57^ This is an important observation, as incomplete sarcolemmal dystrophin coverage is likely to be insufficient to prevent myofiber turnover and correct disease pathology.^55^ (iv) Optimal timing for initiation of treatment. Early initiation of dystrophin restoration therapies may be beneficial to prevent or delay pathological degeneration of muscle. However, for ‘one-and-done’ therapies like microdystrophin gene therapy or CRISPR-Cas9-mediate gene correction, the number of corrected nuclei will be progressively diluted as non-corrected myonuclei are added to myofibers as a consequence of growth and regeneration.

In this study, we assessed the efficacy of exon skipping using PPMO antisense oligonucleotide conjugates when the treatment was initiated at the adult and aged stages. These data support the notion that early treatment may be beneficial in the treatment of DMD.

## Results

### Experimental design

To investigate the efficacy of exon skipping treatment at different ages, *mdx* mice were treated with a peptide-phosphorodiamidate morpholino oligonucleotide (PPMO) Pip6a-PMO (**Figure 1A**), which we have previously shown is effective at inducing skipping of *Dmd* exon 23 and restoring dystrophin protein expression in *mdx* mice.^30,32,40,42,60^ For ‘Adult’ mice (*n*=3), a single 12.5 mg/kg dose of PPMO was administered intravenously at 12 weeks of age and mice sacrificed at 14 weeks of age (**Figure 1B**). For ‘Aged’ mice (*n*=3), three doses of 10 mg/kg PPMO were administered intravenously at 75, 76, and 77 weeks of age and the mice sacrificed at 78 weeks of age. The age of Adult mice was selected as this is a time point after the initial regenerative ‘crisis’ period. The Aged mice were selected based on previous demonstrations of advanced histopathology in this model,^61^ which we reasoned would be more reflective of dystrophic muscle in DMD individuals. Age-, strain-, and sex-matched wild-type (WT, C57BL/10) and dystrophic *mdx* mice were harvested in parallel as controls (all *n*=3). Tibialis anterior (TA) muscles were macrodissected at sacrifice, and subject to a panel of biochemical analyses.

**Figure 1.**
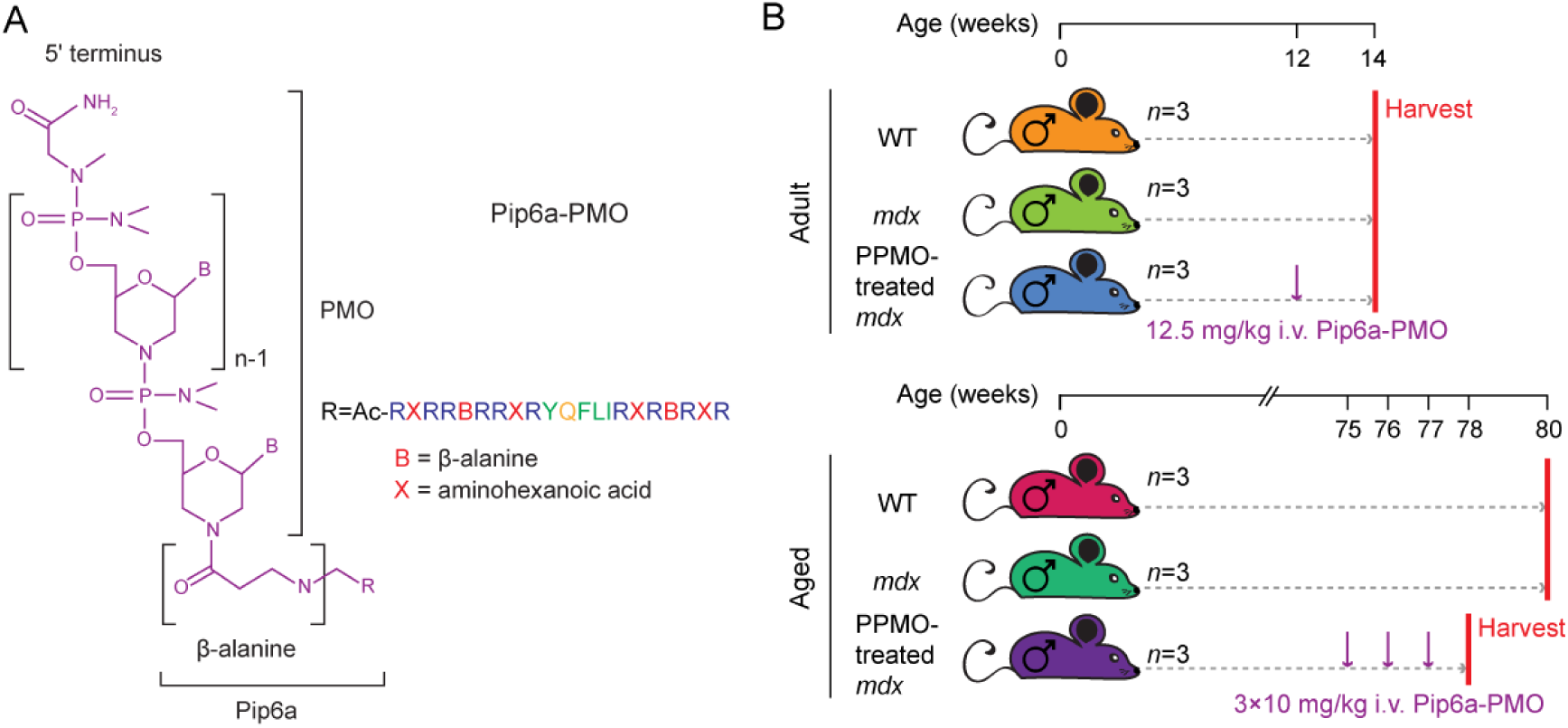
Exon skipping-mediated dystrophin restoration in adult and aged *mdx* mice. (**A**) Structure of the PPMO conjugates, which consist of a Pip6a peptide covalently conjugated to a PMO antisense oligonucleotide designed to skip *Dmd* exon 23. (**B**) Schematic of experimental design. Male *mdx* mice were treated with Pip6a-PMO (PPMO) conjugates with doses and regimens as indicated and harvested at 14 weeks of age for ‘Adult mice’ and at 78 weeks for ‘Aged’ mice. Aged and sex-matched untreated WT and mdx mice were harvested in parallel (all groups *n*=3).

### PPMO treatment at the Adult stage is more effective at restoring dystrophin than treatment at the Aged stage

Mean levels of *Dmd* exon 23 skipping were determined to be ∼79% and ∼44% for the Adult and Aged PPMO-treated *mdx* mice respectively, as determined by RT-qPCR (**Figure 2A**). Restoration of dystrophin protein expression was assessed by western blot, whereby robust re-expression was observed in the Adult PPMO-treated *mdx* mice, while the treatment was much less effective in the Aged mice (**Figure 2B**). Western blot quantification revealed mean dystrophin protein levels were ∼35% and ∼8% of WT levels for the Adult and Aged PPMO-treated *mdx* mice, respectively (**Figure 2C**).

**Figure 2.**
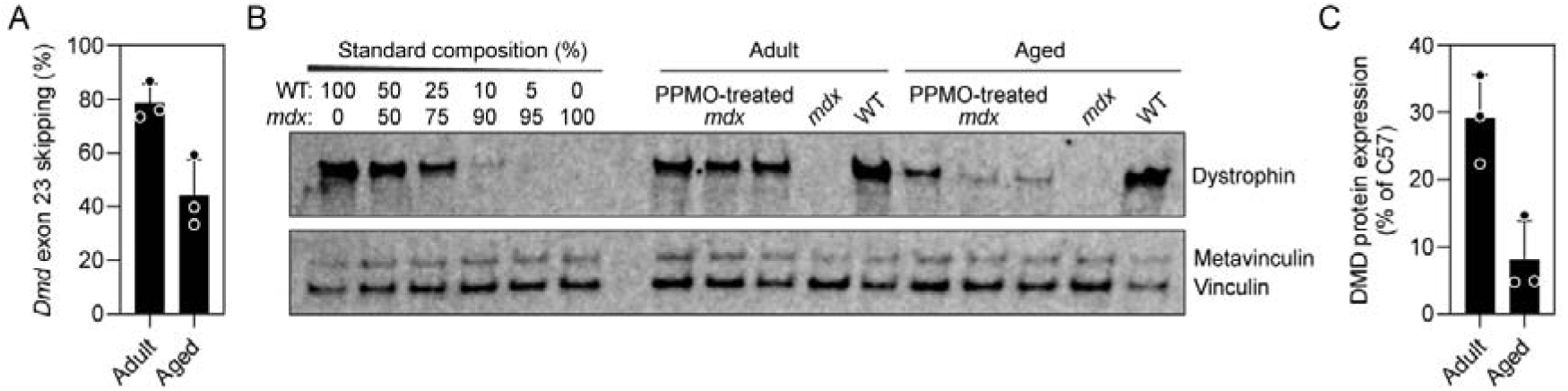
Exon skipping-mediated dystrophin restoration in Adult and Aged *mdx* mice. (**A**) RT-qPCR analysis of *Dmd* exon 23 skipping levels. (**B**) Western blot of dystrophin protein expression for all PPMO-treated *mdx* animals. Vinculin was used as a loading control. (**C**) Quantification of dystrophin protein expression All vales are mean+SD (*n*=3).

Analysis of transverse TA muscle sections revealed a uniform pattern of sarcolemmal dystrophin expression in the PPMO-treated *mdx* animals at both ages (**Figure 3A**). By contrast, untreated *mdx* control animals were dystrophin negative, with the exception of sporadic revertant fibers. Untreated Adult *mdx* mice exhibited characteristic histopathological features of dystrophin absence, including widespread centrally-nucleated fibers and an increase in connective tissue surrounding myofibers (i.e. fibrosis), as assessed by hematoxylin and eosin staining (**Figure 3B**). PPMO-treated *mdx* Adult animals exhibited a reduction in fibrotic damage, while the majority of myofibers were still centrally-nucleated (consistent with them having recently undergone regeneration). We also observed that PPMO-treated Adult myofibers appeared hypertrophic, consistent with previous observations.^55^ Histopathological features were more pronounced in the Aged *mdx* animals, whereby widespread fibrosis was observed, together with irregular fiber sizes and central nucleation. In contrast with the Adult mice, PPMO-treatment did not reverse muscle histopathology in the Aged *mdx* mice (**Figure 3B**).

**Figure 3.**
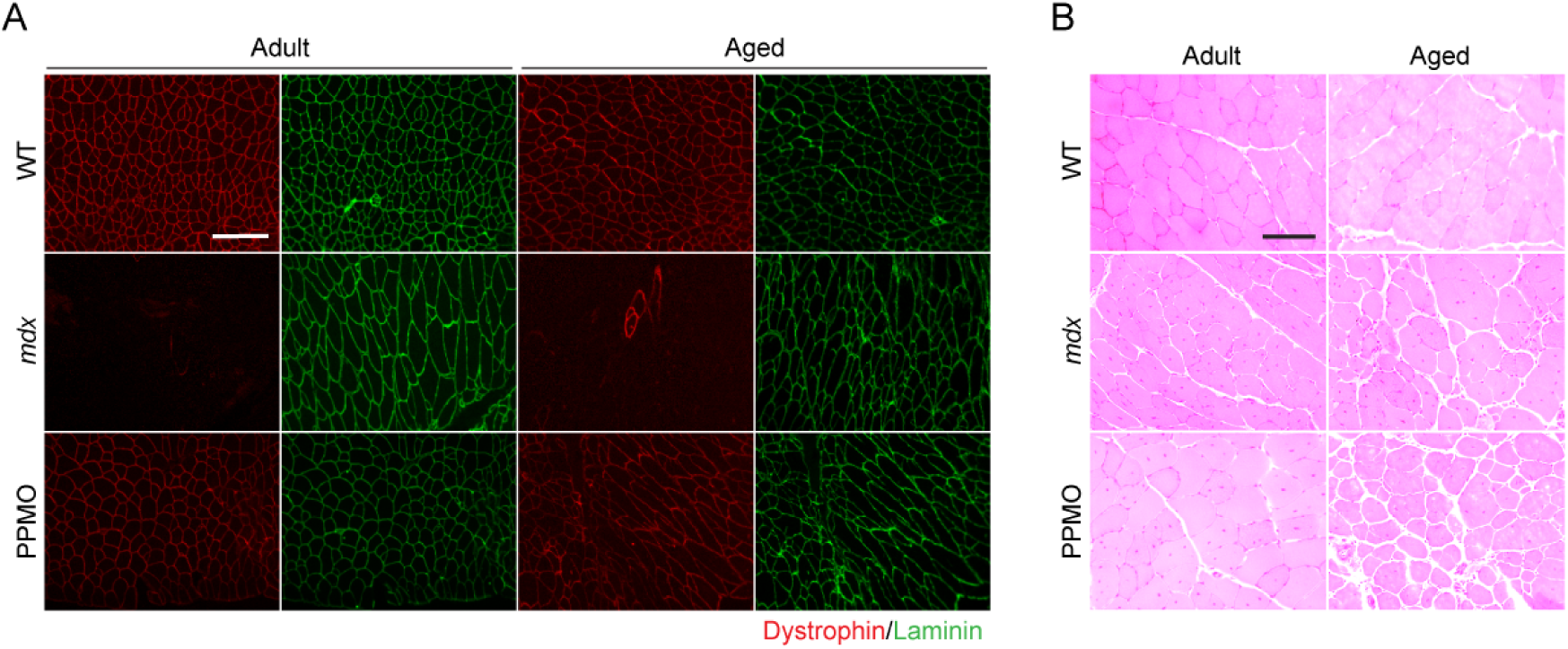
Restoration of sarcolemmal dystrophin expression and histopathological assessment in Adult and Aged *mdx* mice. (**A**) Representative dystrophin immunofluorescence micrographs of TA muscle sections in transverse orientation. Sections were co-stained with laminin to delineate myofiber boundaries. (**B**) H and E staining of TA muscle sections. Images were taken at 20× magnification and scale bars represent 100 µm.

Taken together, these data suggest that the therapeutic effectiveness of PPMO is much greater in Adult animals versus Aged animals, both in terms of the ease with which dystrophin protein can be restored, and the degree to which histopathological changes occurring in dystrophic muscle can be reversed.

### PPMO treatment induces widespread correction of the dystrophic transcriptome at the Adult stage

To better understand the gene expression changes that are occurring in Adult and Aged dystrophic muscle, we performed bulk RNA-Seq analysis in all experimental groups described above (**Figure 1B**). Total RNA samples extracted from TA muscle sections were subjected to Illumina RNA-seq analysis with a target read depth of 50 million reads per sample. Libraries were total RNA, single-end, stranded, and rRNA-depleted. Prior to sequencing, a number of quality control analyses were performed that were indicative of high sample quality, including RNA quantification (pre- and post-rRNA depletion), RNA integrity assessment by Bioanalyzer, and assessment of genomic DNA contamination. (**Figure S1**).

Sequencing reads were processed using a custom in-house pipeline and analysed to identify differentially-expressed genes. Significantly differentially-expressed genes (*P*<0.01) were visualized by heatmap analysis, organized by unsupervised hierarchical clustering (**Figure 2A**). Replicate libraries clustered together for all experimental groups indicative of low inter-replicate variance. Furthermore, WT samples were found to cluster together for both Adult and Aged samples. Conversely, the remaining samples formed the next largest cluster, with the Adult and Aged samples forming sub-clusters (containing a mixture of both *mdx* and PPMO-treated *mdx* samples). Inspection of the heatmap revealed an important qualitative difference between the Adult and Aged samples. Specifically, Adult PPMO-treated animals clustered towards the WT samples, whereas Aged PPMO-treated animals clustered with the *mdx* samples (**Figure 4A**).

**Figure 4.**
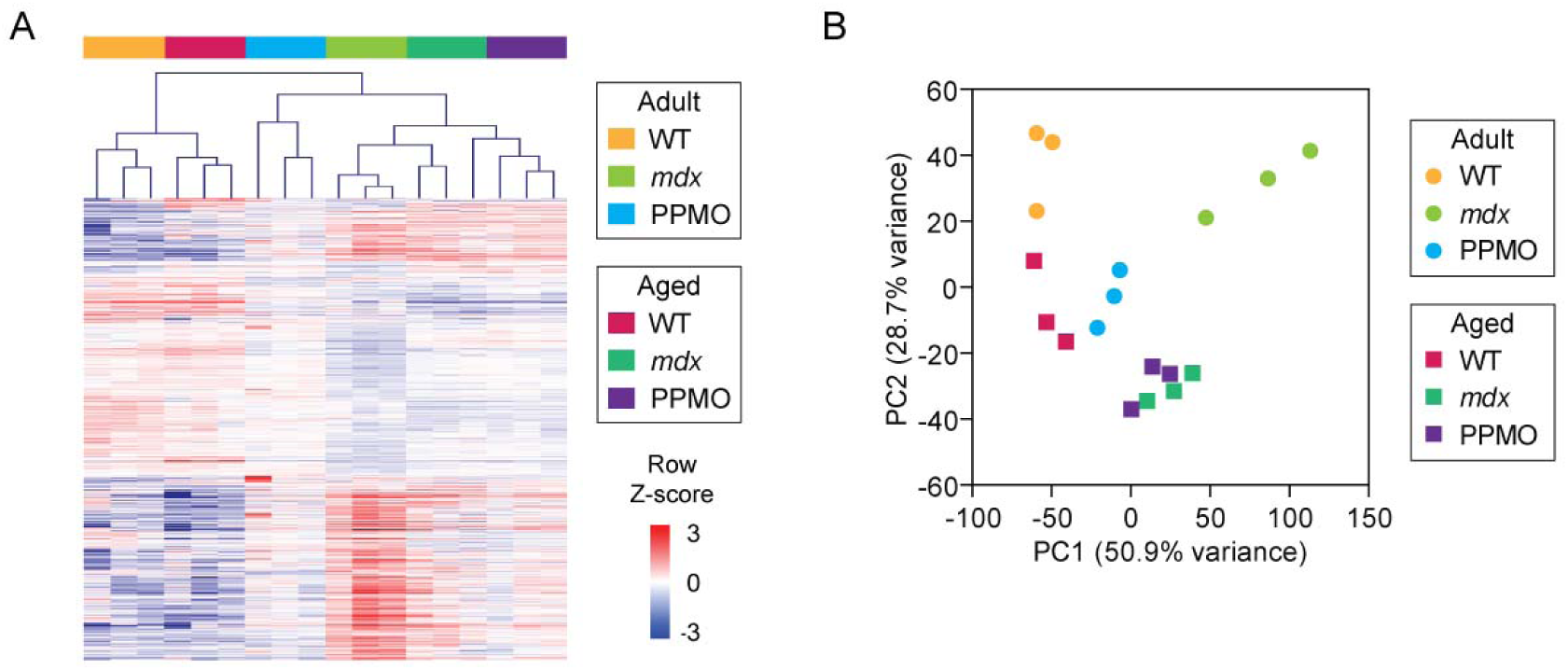
RNA-Seq analysis in Adult and Aged dystrophic muscle. RNA-Seq libraries from WT, *mdx*, and PPMO-treated *mdx* at both Adult and Aged time points were analysed for differential expression. Data are visualized by (**A**) heatmap (unsupervised hierarchical clustering), and (**B**) principal component analysis (PCA) plot.

A similar clustering pattern was observed by principal components analysis (PCA). In general, the difference between dystrophic and WT animals was represented by PC1, which contained 50.9% of the variance (**Figure 4B**). By contrast, PC2 contained 28.7% of the variance, which to some extent reflected the difference between Adult and Aged samples (**Figure 4B**). In Adult animals, PPMO-treated *mdx* animals clustered towards the WT libraries, indicative of a shift (in PC1) from a dystrophic pattern of gene expression towards are more WT-like transcriptome. By contrast, Aged PPMO-treated samples were very closely clustered with the aged-matched *mdx* libraries (**Figure 4B**).

To investigate patterns of differential expression, volcano plots were generated for; (i) *mdx* vs WT, PPMO vs WT, and (iii) PPMO vs *mdx* comparisons for both Adult (**Figure 5A**) and Aged (**Figure 5B**) samples. The greatest number of differentially expressed genes was observed in the Adult *mdx* vs WT comparison (3,946 differentially expressed genes, DGEs). In Adult animals, treatment with PPMO induced profound transcriptomic changes (2,085 DGEs) indicative of a partial reversal in gene expression changes associated with loss of dystrophin. By contrast, there were almost no significant differences between Aged PPMO-treated and *mdx* libraries (7 DGEs).

**Figure 5.**
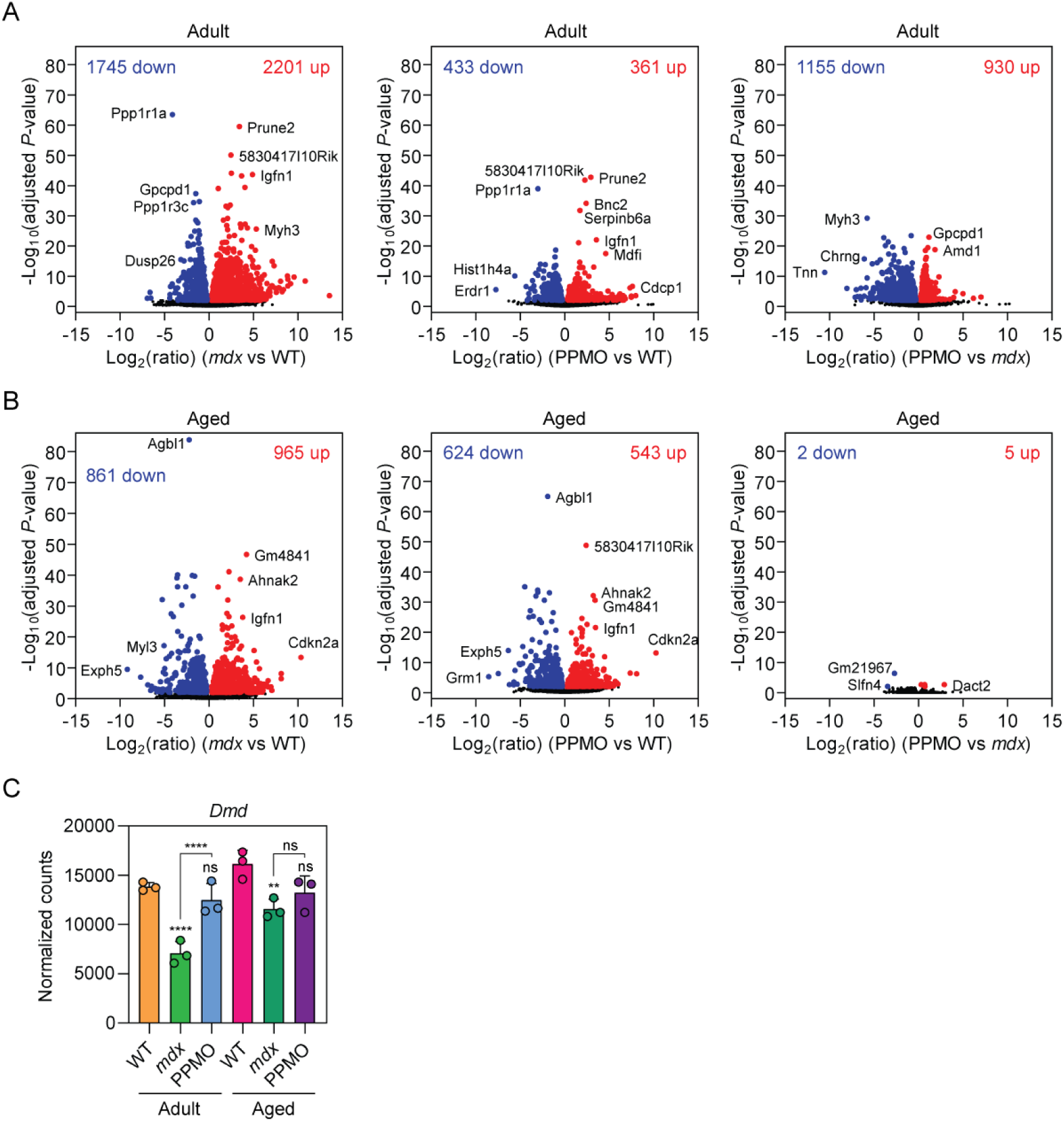
Differential expression analysis in Adult and Aged dystrophic muscle. RNA-Seq libraries from WT, *mdx*, and PPMO-treated *mdx* at both Adult and Aged time points were analysed for differential expression. Volcano plots for each differential expression comparison for (**A**) Adult, and (**B**) Aged mice. (**C**) RNA-Seq normalized counts data for the *Dmd* transcript for all libraries. Statistical significance was assessed using DESeq2 and Benjamini-Hochberg-adjusted *P*-values indicated. Statistical comparisons are relative to the Adult WT group unless otherwise indicated, ****adjusted-*P*<0.0001, ns=not significant.

A reduction in *Dmd* mRNA levels is a characteristic feature in *mdx* mice,^40,62,63^ that has been attributed to a transcriptional mechanism.^64^ Accordingly, in Adult animals, *Dmd* expression was reduced by ∼50%, and was restored to near-WT levels by PPMO treatment (**Figure 5C**). By contrast, in Aged animals, *Dmd* levels were reduced by ∼30% in *mdx* animals and were not restored by PPMO treatment (**Figure 5C**).

Together, these data are indicative of widespread transcriptional perturbations in dystrophic muscle, which are more complex at the Adult time point. PPMO-treatment results in a shift in the transcriptional profile of Adult dystrophic muscle towards that of WT muscle. By contrast, PPMO-treatment of Aged muscle had negligible influence on the dystrophic transcriptome.

### PPMO treatment reverses *Dmd* transcript imbalance in Adult dystrophic muscle

*mdx* animals are known to exhibit a transcript imbalance effect, whereby dystrophic muscle contains greater quantities of the 5L end of the *Dmd* transcript and reduced levels of the 3L end, compared to WT muscle.^65,66^ This phenomenon was observed in both Adult and Aged *mdx* mice when RNA-Seq data were analysed for differential exon usage using DEXSeq^67^ (**Figure 6**). Treatment with PPMO reversed the transcript imbalance effect in Adult *mdx* mice (**Figure 6A,B**), but not in Aged *mdx* mice (**Figure 6C,D**). Notably, it has previously been reported that transcript imbalance is not corrected by exon skipping therapy, although the threshold of dystrophin restoration required to correct this phenomenon may not have been achieved in these studies.^66,68^

**Figure 6.**
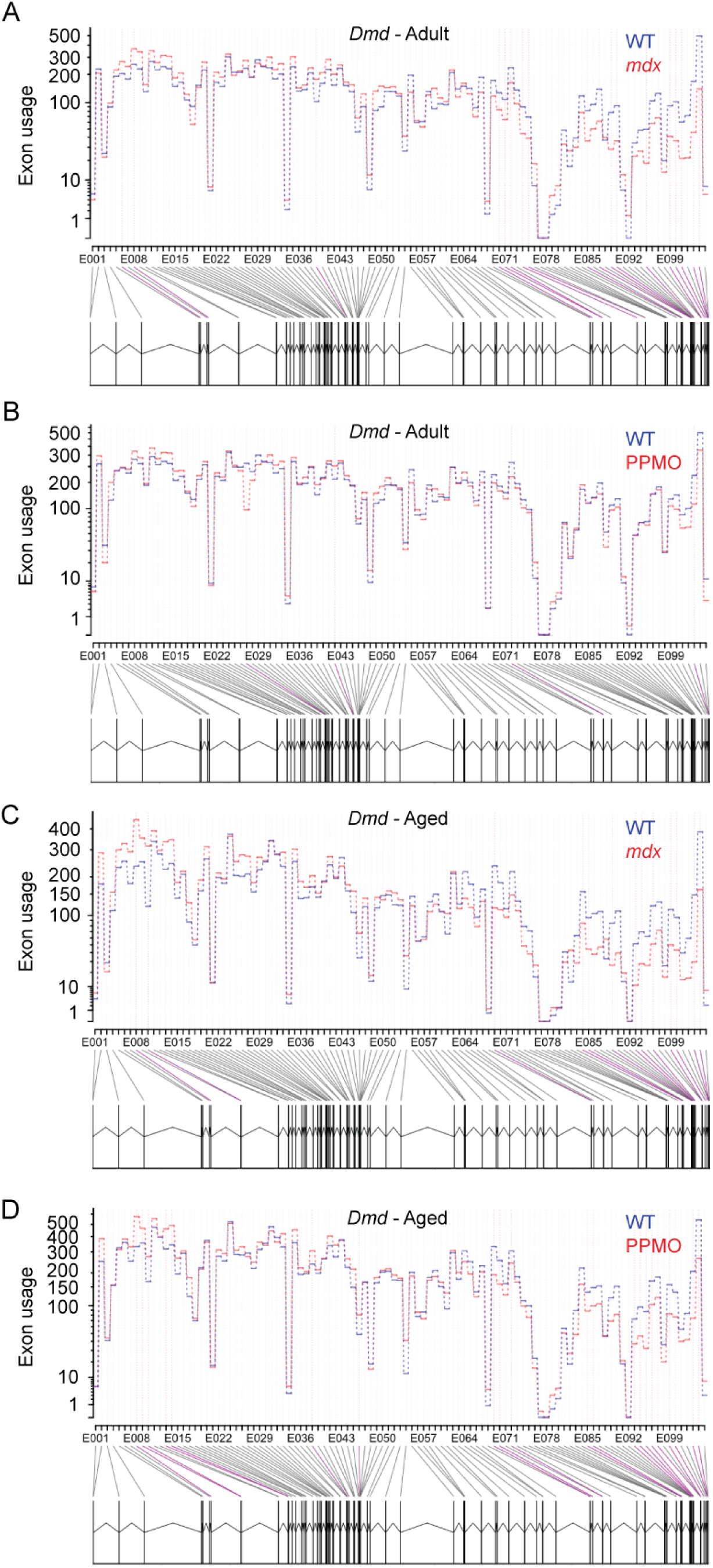
PPMO treatment restores Dmd transcript imbalance in Adult but not Aged *mdx* mice. Differential exon usage for the *Dmd* transcript was analysed using and visualised using RNA-Seq data for four comparisons: (**A**) Adult *mdx* vs WT, (**B**) Adult PPMO vs WT, (**C**) Aged *mdx* vs WT, and (**D**) Aged PPMO vs WT. Exons showing statistically significant differential usage are highlighted with purple lines.

### Differential gene expression signatures in dystrophic muscle

To characterise the biological meaning of gene expression changes occurring in dystrophic muscle, we performed gene list enrichment analysis and semantic space reduction DGE lists for the *mdx* vs WT at the Adult (**Figure S2**) and Aged (**Figure S3**) stages. These analyses identified some expected perturbed pathways, including calcium ion binding, electron transfer activity, extracellular matrix organization, immune system process, reactive oxygen species metabolic process, and regulation of nitric oxide metabolic process, but also many other GO terms that were mostly generic and not otherwise informative. By contrast, manual inspection of gene expression changes revealed gene expression signatures that are consistent with known aspects of dystrophic pathology. *Myh3* (embryonic myosin heavy chain) was upregulated at both Adult and Aged time points, although the magnitude of upregulation was much higher at the Adult stage. Interestingly, PPMO-treatment restored *Myh3* expression to WT levels in both cases, indicative of a stabilization of muscle turnover, even in Aged muscle (**Figure S4A**). Concordantly, *Pax7* expression was highly increased in *mdx* muscle at the Adult stage only. This corresponds with the high level of muscle regeneration occurring at this age, with a concomitant expansion of the satellite (stem) cell pool (**Figure S4B**). The senescence markers *Cdkn1a* (p21) and *Cdkn2a* (p16) were strongly upregulated in *mdx* muscle, with *Cdkn1a* more prominent at the Adult stage and *Cdkn2a* more pronounced at the Aged stage. This is indicative of accumulated cell stress, which may contribute to a functionally compromised regenerative niche, especially in aged dystrophic muscle (**Figure S4C**). Neuronal nitric oxide synthase protein mislocalization is a characteristic feature of dystrophic muscle. However, we also observed downregulation of *Nos1* transcripts, which was more apparent at the Aged time point (**Figure S4D**). *Nos3* was similarly downregulated in dystrophic muscle. *Vegfa* and its receptor *Kdr* were also downregulated in dystrophic muscle. These data are consistent with gene expression changes that promote functional ischemia in *mdx* mouse muscle (**Figure S4D**).

Numerous transcripts encoding proteins involved in reactive oxygen species handling were downregulated in dystrophic muscle, including *Egln1*, *Sod2*, *Ndufs1*, *Prdx3* (**Figure S5**). Additionally, *Ucp3*, *Epas1*, *Cox4i1*, and *Txnrd1* were similarly downregulated, with Aged animals exhibiting a further decrease (**Figure S5**). The antioxidant *Gpx3* was highly elevated in Aged WT animals, but did show a corresponding increase in Aged *mdx* animals. These findings are consistent with a failure of dystrophin-deficient muscle to effectively manage ROS production. *Ppargc1* (PGC1-α) was downregulated in *mdx* tissues at both ages (**Figure S5**), consistent with a suppression of mitochondrial biogenesis and oxidative metabolism, which may further contribute to the oxidative stress phenotype in dystrophic muscle.

Persistent inflammation is a characteristic feature of dystrophic muscle, which was reflected in the RNA-Seq gene expression patterns (**Figure S6A**). Specifically, leukocyte (*Ptpr*c), T cell (*Cd3e* and *Cd4*), and macrophage/monocytes (*Cd68* and *Itgam*) markers were upregulated in dystrophic muscle. The dendritic cell marker (*Itgax*) was strongly upregulated at the Adult stage, possibly associated with the peak of active regeneration observed at this time point. Transcript levels for cytokines and chemokines (e.g. *Tnf*, *Il1b*, *Il6*, *Ccl2*) were undetectable or not-significantly changed between libraries (data not shown). Components of the non-canonical NF-κB pathway (*Relb* and *Nfkb2*) were significantly increased in *mdx* muscle (**Figure S6B**), consistent with a chronic, low-grade inflammatory or inflammaging-like state.

Markers of fibrosis were upregulated in *mdx* muscle including *Col1a1*, *Col3a1*, *Col6a1*, *Fn1*, *Ctgf*, *Timp1*, *Mmp2*, and *Ctgf* (**Figure S7**). Interestingly, these transcripts exhibited very similar patterns of expression, with a higher degree of upregulation in Adult *mdx* muscle than in Aged *mdx* muscle. This pattern of expression may reflect the shift from the period of active regeneration observed in the Adult mdx mice (with associated intense ECM remodelling) to a period of more stable period of fibrotic decline. We have previously observed an elevation in COL3A1 protein expression only in aged dystrophic mice (*mdx* and mdx52 *strains*) by mass-spectrometry-based proteomics analysis, highlighting a divergence between gene expression and tissue protein levels. Notably, *Col1a1* and *Col3a1* were reduced in Aged WT muscle, further highlighting that differences in the expression of these transcripts in dystrophic muscle substantially diverge from the pattern expected during the course of normal aging. *Tgfb1* was only significantly upregulated at the adult stage. Key markers of adipo-degeneration were unchanged across all libraries (i.e. *Pparg*, *Cepba*, *Fabp4*, and *Plin1*, data not shown) consistent with minimal pathology of this type observed in *mdx* mice.

We next sought to investigate possible reasons for impaired re-expression of dystrophin expression in Aged *mdx* muscle following PPMO treatment. Firstly, we investigated the possibility that a global downregulation of translation may underlie the impairment in dystrophin re-expression. The integrated stress response (ISR) is a pathway that controls the response to cellular stresses via the phosphorylation of eIF2α (eukaryotic initiation factor 2 alpha), which results in a global downregulation of translation while promoting selective activation of stress-responsive genes. Transcript levels for key ISR components were unchanged across all libraries (*Atf4*, *Ddit3*, *Eif2ak1*, *Eif2ak2*, *Eif2ak3*, *Eif2ak4*, and *Eif2s1, Ppp1r15a* (GADD34); data not shown). Similarly, transcript levels for multiple downstream ATF4 targets were either unchanged (*Asns*, *Trib3*, and *Psat1*, *Ppp1r15a* [GADD34] data not shown), suggesting that the ISR is not engaged in this context. Similarly, transcripts associated with the unfolded protein response (UPR) arm of the ISR were similarly unchanged (*Hspa5*, *Hsp90b1*, *Pdia3*, and *Calr*; data not shown). While ISR activation has been reported in DMD muscle and dystrophic animal models,^69,70^ the data presented herein argue against the ISR involvement as a mechanism underlying impaired dystrophin re-expression in Aged *mdx* muscle.

The AKT/PI3K/mTOR signalling axis is an important regulator of protein synthesis in muscle. Analysis of transcript levels associated with this pathway points to a mixed picture, with no significant changes in *Ddit4*, *Tsc1*, *Slc1a5*, *Atg13*, *Trim63*, *Eif4g1*, *Mtor*, *Map1lc3b*, *Tsc2*, *Pik3r1*, or *Eif4ebp1* (data not shown). *Akt1* and *Myc* were elevated in Adult *mdx* muscle, suggestive of a pro-growth phenotype consistent with muscle regeneration occurring at this age (**Figure S8**). However, *Slc38a2*, *Pik3ca*, *Prkaa2*, *Rheb*, *Insr*, *Rps6kb1*, *Rictor*, *Eif4e* and *Fxbo32* were down-regulated in Adult *mdx* muscle indicative of potentially reduced AKT/PI3K/mTOR pathway activity (**Figure S8**). *Igf1r* was down-regulated in Aged *mdx* muscle (**Figure S8**). Notably, previous proteomics analyses from our group identified the PI3K/AKT pathway as being activated in dystrophic muscle at 8- and 16-weeks, and repressed in 80-week-old mice.^43^ These data suggest that attenuated mTOR signalling may contribute to impaired dystrophin re-expression in Aged dystrophic muscle.

### miR-31 upregulation is further enhanced in Aged dystrophic muscle

Previous work has shown that microRNAs (miRNAs) can repress expression of dystrophin expression, which can limit the effectiveness of exon skipping therapies in *mdx* mice treated at the adult stage.^71,72^

The most well-characterized example, miR-31-5p, is highly upregulated in dystrophic muscle.^40–42,71,73^ To investigate the role of miRNAs in stage-specific PPMO-treatment efficacy, we measured the expression of miR-31-5p in all treatment groups. Two additional miRNAs, miR-34c-5p and miR-206-3p were included as they are known to be differentially expressed in dystrophic muscle but are not expected to regulate dystrophin expression *per se*.^40–42,73,74^ miR-31-5p was found to be highly elevated in Adult *mdx* muscle (∼60-fold, *P*<0.05) and was not corrected by PPMO treatment (**Figure 7A**). miR-31-5p upregulation was more pronounced in Aged *mdx* muscle (∼120-fold, *P*<0.001) with PPMO-treatment leading to an even greater increase (∼186-fold relative to WT, *P*<0.0001, **Figure 7A**). By contrast, miR-34c-5p was upregulated in *mdx* muscle at the Adult stage only (∼6.6-fold, *P*<0.01, **Figure 7B**) consistent with previous reports,^40–42,73^ and restored to WT levels by PPMO treatment. miR-206-3p was increased by similar magnitudes (∼6-fold, *P*<0.01, **Figure 7C**) in dystrophic muscle at both time points. These data suggest that an intensification in miR-31-5p expression in Aged dystrophic muscle is a possible mechanistic explanation for the limited efficacy of PPMO-mediated exon skipping therapy at this age.

**Figure 7.**
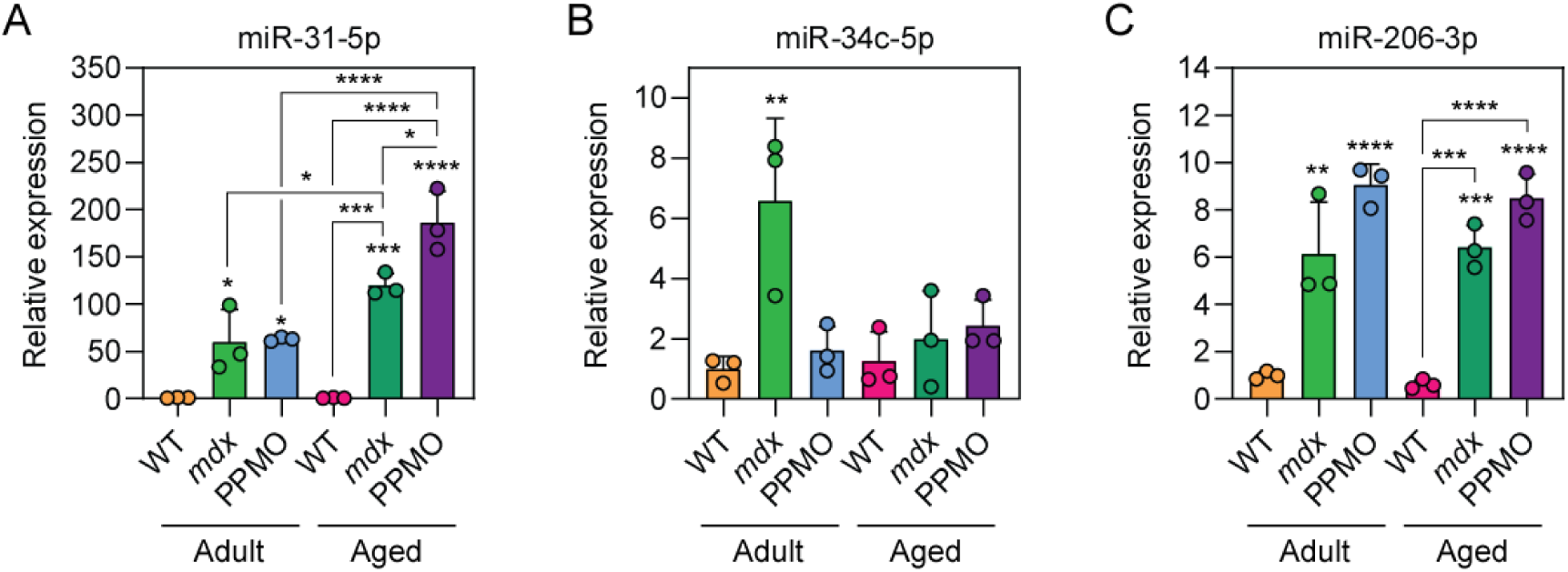
Upregulated miR-31 expression in Adult dystrophic muscle is further increased in Aged dystrophic muscle. RNA samples from WT, *mdx*, and PPMO-treated *mdx* mice at Adult and Aged time points were analysed by RT-qPCR to determine differential miRNA expression for (**A**) miR-31-5p, (**B**) miR-34c-5p, and (**C**) miR-206-3p (normalized to miR-16-5p). All values are mean+SD (*n*=3). Statistical significance was tested by one-way ANOVA with Tukey’s *post hoc test*. Statistical comparisons are relative to the Adult WT group unless otherwise indicated, \**P*<0.05, \*\**P*<0.01, \*\*\**P*<0.001, \*\*\*\**P*<0.0001.

## Materials and Methods

### Animals

Male wild-type C57 (C57BL/10) and dystrophic *mdx* (C57BL/10ScSn-Dmd^mdx^/J) mice (both *n*=3) were harvested at either at two ages; adult (14 week) and aged (80 week). PPMO conjugates consisting of a PMO (5’-GGCCAAACCTCGGCTTACCTGAAAT, Gene Tools) conjugated by amide linkage to Pip6a (Ac-RXRRBRRXRYQFLIRXRBRXRB-OH, X=aminohexanoyl, B=β-Alanine) were prepared in sterile saline as described previously ^30^. All animals were injected via the tail vein under isoflurane anaesthesia. All mice were sacrificed by escalating CO_2_ concentration, and death confirmed by cervical dislocation. Tibialis anterior (TA) muscles were macrodissected immediately post mortem, mounted onto cork discs using OTC media and flash frozen in liquid nitrogen-cooled isopentane. Animal experimentation was carried out in accordance with the Animals (Scientific Procedures) Act 1986 and all procedures approved by the UK home office.

### RNA Preparation

RNA was extracted from 10 mg of TA tissue using the Maxwell 16 RSC instrument with the (Maxwell RSC Simply RNA Tissue kit) (both Promega), which includes a rigorous DNase step. RNA concentration and quality were assessed by Nanodrop spectrophotometry to measure absorbance at 260 nm and the 260/280 and 260/230 ratios. RNA integrity assessed by Bioanalyzer (Eukaryote RNA Pico chip) (Agilent). Contamination of RNA samples with genomic DNA was assessed by qPCR (with or without reverse transcription) using a TaqMan assay, which amplifies *Actb* mRNA/genomic DNA (Mm00607939_s1). RNA samples (1 µg each) were depleted of ribosomal RNA using the NEBNext rRNA Depletion Kit (Human/Mouse/Rat) (NEB) according to manufacturer’s instructions with minor modifications (reaction volumes were scaled to accommodate sample RNA concentrations). rRNA-depleted RNA samples were eluted in 10 µl of nuclease-free water and 8 µl retained for further analysis. The concentrations of rRNA-depleted RNA samples were determined by RNA fluorometry using the Quantus instrument and appropriate reagents (QunatiFluor RNA System) (both Promega). RNA quality control is summarised in **Figure S1**.

### RNA-Seq

RNA-Seq was performed as a service at Karolinsk Institutet on the Illumina NextSeq 500 platform. Sequencing reads were processed using a custom in-house pipeline. In brief, the suitability of FASTQ files for analyses was using FastQC (v0.10.1) and MultiQC (v0.7).^82^ Reads were trimmed using Trim_Galore (v0.3.1) and aligned to the mouse genome (GRCm38) using HISAT2 (v.2.1.0).^83^ Alignment files were processed using Samtools (v1.3).^84^ Reads were counted using HTSeq (v0.9.1) against the v96 GTF file obtained from Ensembl.^85^ Differential expression was assessed using DESeq2 (v1.28.1). Differential exon expression usage was assessed using DEXSeq (v3.8).^67^ Gene ontology analysis was performed using a combination of g:Profiler and REVIGO.^86,87^ Raw RNA-Seq data are deposited in NCBI SRA under BioProject accession: PRJNA1384431. Counts data are provided in **Figure S1**.

### Exon Skipping RT-qPCR

Exon skipping was confirmed by RT-qPCR to measure the levels of *Dmd* transcripts lacking exon 23. Briefly, reverse transcription was performed on 1 µg total RNA using the High Capacity cDNA Kit (Thermo Fisher Scientific, Rugby, UK) according to manufacturer’s instructions. Levels of *Dmd* Δ23 transcripts were determined using primer/probe assays (Integrated DNA Technologies, Leuven, Belgium) spanning the exon 23-24 boundary (to measure unskipped *Dmd* transcripts) and spanning the exon 20-21 boundary (to measure total *Dmd* transcripts) (**Table S1**). Probes were HEX and FAM labelled respectively and run in singleplex format. qPCR was performed on a StepOne Plus real-time PCR Thermocycler using TaqMan Gene Expression Master Mix (both Thermo Fisher Scientific) and 25 ng cDNA template was added per reaction. The percentage of *Dmd* exon 23 skipping was determined by calculating (1 – the ratio of unskipped:total *Dmd* transcripts) × 100%.

### Dystrophin Western Blot

For dystrophin protein quantification, 8 µm cryosections (∼80) were prepared from the mid-belly of TA muscle and samples lysed in Lysis Buffer (50 mM Tris pH 8, 150 mM NaCl, 1% NP-40, 0.5% sodium deoxycholate, 10% sodium dodecyl sulphate, and protease inhibitors). Protein lysates were incubated at 100°C for 3 minutes and then centrifuged at 14,000 *g* for 10 minutes at 4°C to pellet debris and supernatants transferred to fresh tubes. Equal amounts of protein lysate (20 µg) were separated on a 3-8% Tris-Acetate gel (Life Technologies), electrotransferred to a Polyvinylidene fluoride (PVDF) membrane and probed with monoclonal anti-dystrophin (NCL-DYS1, Leica Biosystems, Lincoln, NE) and anti-vinculin (hVIN-1 V9131, Sigma, Dorset, UK) primary antibodies). Secondary antibody fluorescence was quantified using the Odyssey imaging system (IRDye 800CW goat anti-mouse, LiCOR Biosystems). To quantify dystrophin expression, the dystrophin to vinculin ratio for each PPMO-treated sample was compared to a dilution series of C57Bl/10 lysate (diluted in *mdx* lysate to maintain consistent vinculin expression levels between standards) as described previously.^44^

### Histology and Immunofluorescence

8 µm sections of TA muscles were analysed at the Histology Core, Kennedy Institute of Rheumatology, University of Oxford.

### miRNA RT-qPCR

miRNAs were quantified using the small RNA TaqMan RT-qPCR method using miRNA-specific stem loop reverse transcription primers. Details of miRNA assays are listed in **Table S2**. Reverse transcription was performed using the TaqMan MicroRNA Reverse Transcription Kit (Thermo Fisher Scientific) with and 20 ng of input total RNA per reaction. cDNA was amplified using a StepOne Plus real-time PCR thermocycler with TaqMan Gene Expression Master Mix (both Thermo Fisher Scientific) using universal cycling conditions: 95°C for 10 minutes, followed by 40 cycles of 95°C for 15 seconds and 60°C for 1 minute. All samples were analysed in duplicate. Relative quantification was performed using the Pfaffl method,^88^ and miRNA of interest expression normalized to miR-16-5p.

### Statistics

Statistical analysis was performed in GraphPad Prism (v10.2.3) (GraphPad Software Inc., San Diego, California, USA) using an ordinary one-way analysis of variance (ANOVA) with Tukey’s *post hoc* test for inter-group comparisons.

## Discussion

Advanced therapeutics that can restore dystrophin protein expression in the muscles of DMD patients are now reaching the clinic, with multiple approvals in the US for exon skipping and microdystrophin gene replacement approaches. Next-generation exon skipping approaches are on the horizon, which will likely offer improved dystrophin restoration potential. These developments bring into focus the question of treatment timing.

The data presented herein support the idea that exon skipping treatments should be initiated early, in order to maximise therapeutic benefit. A single intravenous dose of a PPMO conjugate resulted in high levels of *Dmd* exon23 skipping and dystrophin protein restoration (∼35% of WT levels) after only 2 weeks of treatment in Adult mice (**Figure 2**). This was associated with a reversal of histopathological changes (**Figure 3B**) and a major shift in the dystrophic muscle transcriptome (**Figure 4,5**), indicative of therapeutic success. By contrast, three intravenous doses (2.4-times the total dose administered at the Adult stage) resulted in more modest dystrophin restoration (∼8%) (**Figure 2**), with negligible histopathological or transcriptomic restoration (**Figure 3,4,5**). Together, these data show that the timing of exon skipping treatment is an important determinant of treatment efficacy.

The relative lack of therapeutic effectiveness may be a consequence of a number of factors. Firstly, reduced treatment efficacy may be the result of an impairment in drug delivery. The muscles of Aged *mdx* mice exhibit a greater degree of fibro-fatty degeneration when compared to the adult stage, which resembles the pathology observed in DMD patients.^61^ However, inefficient delivery is insufficient to explain all of the data. Specifically, the levels of PPMO-mediated protein restoration were ∼4 times lower in Aged versus Adult treated animals, while the level of PPMO-mediated exon skipping was only ∼1.8 less (**Figure 2**). Indeed, the level of *Dmd* exon23 skipping in the Aged PPMO-treated animals was still relatively high (at 44%), indicative of successful drug delivery and target engagement (although still at a reduced level relative to the Adult PPMO-treated mice) (**Figure 2**).

Secondly, it is possible that dystrophin restoration in the degenerated dystrophic muscle environment may be functionally inconsequential. In this case, dystrophin restoration may be successful (at least to some extent) in the remaining myofibers, but insufficient to reverse fibro-fatty muscle degeneration (i.e. muscle quality). The lack of transcriptomic response to PPMO treatment in Aged *mdx* animals supports this notion (**Figure 4,5**).

Lastly, it is possible that the Aged dystrophic environment is inherently resistant to dystrophin restoration as a consequence of molecular and cellular pathologies. Such concepts have been reported previously, whereby dystrophic muscle is similarly refractory to therapeutic correction, including impaired AAV transgene expression in the post-regeneration environment,^75^ direct oxidative damage to the *Dmd* transcript itself,^76^ and suppression of dystrophin protein expression via miRNAs.^71,72^ Indeed, tissue miRNA analysis showed that miR-31-5p is elevated in Adult dystrophic muscle, and elevated even further in Aged dystrophic muscle (**Figure 7**). It is likely that a confluence of factors (e.g. multiple miRNAs, RNA binding proteins, oxidative stress, gene expression changes, and cellular senescence, AKT/PI3K/mTOR axis perturbation) may act synergistically to limit exon skipping treatment effectiveness in Aged dystrophic muscle.

Importantly, early therapeutic intervention in the case of spinal muscular atrophy (SMA) has proven critical for therapeutic success.^77^ Whether such a benefit of early intervention in DMD patients remains to be demonstrated. Furthermore, an ongoing challenge in the treatment/management of DMD is the problem of diagnostic delay (the average age at diagnosis is 5 years, with an average time from first signs to diagnostic confirmation of 2.2 years),^78^ which hampers efforts towards early initiation of therapy. This issue highlights a need for newborn screening programs for early diagnosis of DMD.^79^ The approval of the Elevidys microdystrophin gene therapy (which is potentially applicable to the majority of DMD patients) strengthens the case for newborn screening, although there are important safety concerns related to this treatment following several instances of fatal acute liver failure in non-ambulatory DMD patients.^28,80^

Exon skipping therapies may also be implemented as part of a combination therapy. In this manner, exon skipping therapy could be initiated in young DMD boys with the aim of maintaining their muscle function until such an age that treatment with a gene therapy may be safe and appropriate. Indeed, preclinical evidence is supportive of a synergistic effect of combined PMO-mediated exon skipping with either AAV-U7-mediated exon skipping or AAV-microdystrophin gene therapy.^81^

This study adds to the discourse concerning the appropriate time to start exon skipping therapy in DMD patients. Evidence from the preclinical data presented herein supports the notion of initiating exon skipping therapy at an early age, before the establishment of advanced tissue pathology.

## Acknowledgements

This work was supported by a grant from the UK Medical Research Council (MR/X008029/1) awarded to MJAW and TCR. SS was funded by an MRC Confidence-in-Concept grant awarded to MJAW and TCR. TLEvW was supported by a doctoral studentship from Muscular Dystrophy UK (MDUK) awarded to MJAW and TCR.

## Author Contributions

TCR, SEA, and MJAW conceived the study. TCR, SEA, and MJAW supervised the work. SS, JH, TLEvW, AMLC-S, and KK performed experimentation. TCR wrote the first draft of the manuscript. All authors contributed to and approved the final version of the manuscript.

## Data Availability Statement

Raw data have been deposited in NCBI SRA under BioProject accession: PRJNA1384431. Processed counts data are included as a supplemental file. All other data are contained in the manuscript.

