## Supplemental Information for "Timing matters: exon skipping therapy is most effective when initiated early in a mouse model of Duchenne muscular dystrophy"

**Table S1**

**RT-qPCR assays used in this study.**

All sequence are 5′ to 3′.

| | | Sequence |
|---|---|---|
| *Dmd* exon 20-21 | FWD | AGATGACAACTACTGCCGAA |
| | REV | GAAGAGCTGACAATCTGTTGAC |
| | PROBE | AGTCTACCACCCTATCAGAGCCAACA |
| *Dmd* exon 23-24 | FWD | GAAACTTTCCTCCCAGTTGGT |
| | REV | CAGGCCATTCCTCTTTCAGG |
| | PROBE | TCAACTTCAGCCATCCATTTCTGTAAGGT |

**Table S2**

**List of Small NA TaqMan RT-qPCR assays used in this study.**

All assays were obtained from Thermo Fisher Scientific.

| Target | Product ID |
|---|---|
| mmu-miR-16-5p | 000391 |
| mmu-miR-31-5p | 000185 |
| mmu-miR-34c-5p | 000428 |
| mmu-miR-206-3p | 000510 |

A

| | Sample ID | Total RNA ng/µl | Total RNA A260/A280 | Total RNA A260/A230 | Total RNA RIN | rRNA minus ng/µl |
|---|---|---|---|---|---|---|
| Adult (14 weeks) | C57 1 | 77.2 | 2.11 | 1.85 | 9.6 | 16 |
| | C57 2 | 72.4 | 2.09 | 1.79 | 8.4 | 10 |
| | C57 3 | 63.9 | 2.08 | 1.77 | 9.2 | 9 |
| | *mdx* 1 | 154.3 | 2.07 | 1.97 | 7.8 | 8.8 |
| | *mdx* 2 | 65.7 | 2.03 | 1.71 | 6.7 | 8.7 |
| | *mdx* 3 | 84.7 | 2.08 | 1.73 | 7.6 | 15 |
| | PPMO 1 | 73.3 | 2.06 | 1.79 | 9.4 | 11 |
| | PPMO 2 | 76 | 2.05 | 1.62 | 9.2 | 13 |
| | PPMO 3 | 78.3 | 2.07 | 1.79 | 9.2 | 9.8 |
| Aged (78-80 weeks) | C57 1 | 62.2 | 2.09 | 1.77 | 9.2 | 15 |
| | C57 2 | 45.4 | 2.01 | 1.58 | 8.9 | 12 |
| | C57 3 | 46.9 | 2.07 | 1.71 | 8.7 | 7.4 |
| | *mdx* 1 | 70.5 | 2.06 | 1.77 | 8.1 | 11 |
| | *mdx* 2 | 61 | 2.08 | 1.82 | 7.9 | 14 |
| | *mdx* 3 | 51.9 | 2.07 | 1.72 | 8.4 | 14 |
| | PPMO 1 | 60.8 | 2.03 | 1.53 | 8.7 | 12 |
| | PPMO 2 | 46.5 | 2.01 | 1.63 | 8.4 | 7.5 |
| | PPMO 3 | 77.5 | 2.06 | 1.71 | 9 | 4.8 |

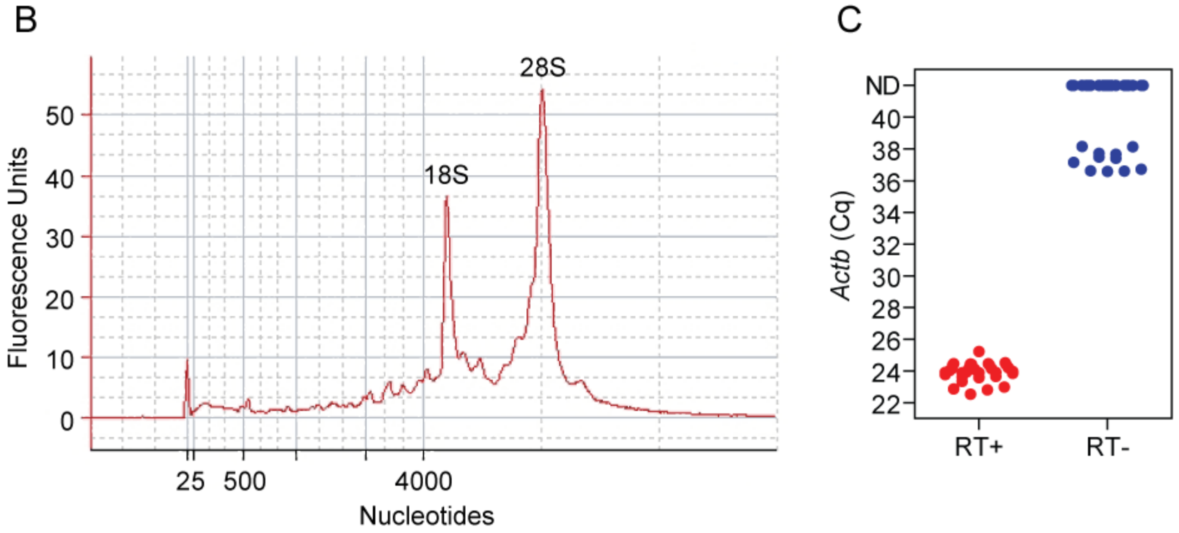

**Figure S1**

**RNA-Seq quality control analyses.**

(**A**) Summary of sample assessments of RNA concentration, quality, and integrity by UV spectrophotometry, bioanalyzer, and fluorimetry (post rRNA depletion). (**B**) Typical bioanalyzer trace indicating intact RNA samples with prominent rRNA peaks (pre rRNA removal). (**C**) RNA samples were assessed by genomic DNA contamination by measurement of *Actb* mRNA expression by RT-qPCR. This assay is designed in such a way that it has the potential to amplify both mRNA and its encoding DNA. Amplification in a RT- control sample would therefore be indicative of genomic DNA contamination. *Actb* levels in the RT- samples were either note detected (ND) or in the single copy range (Ct >35).

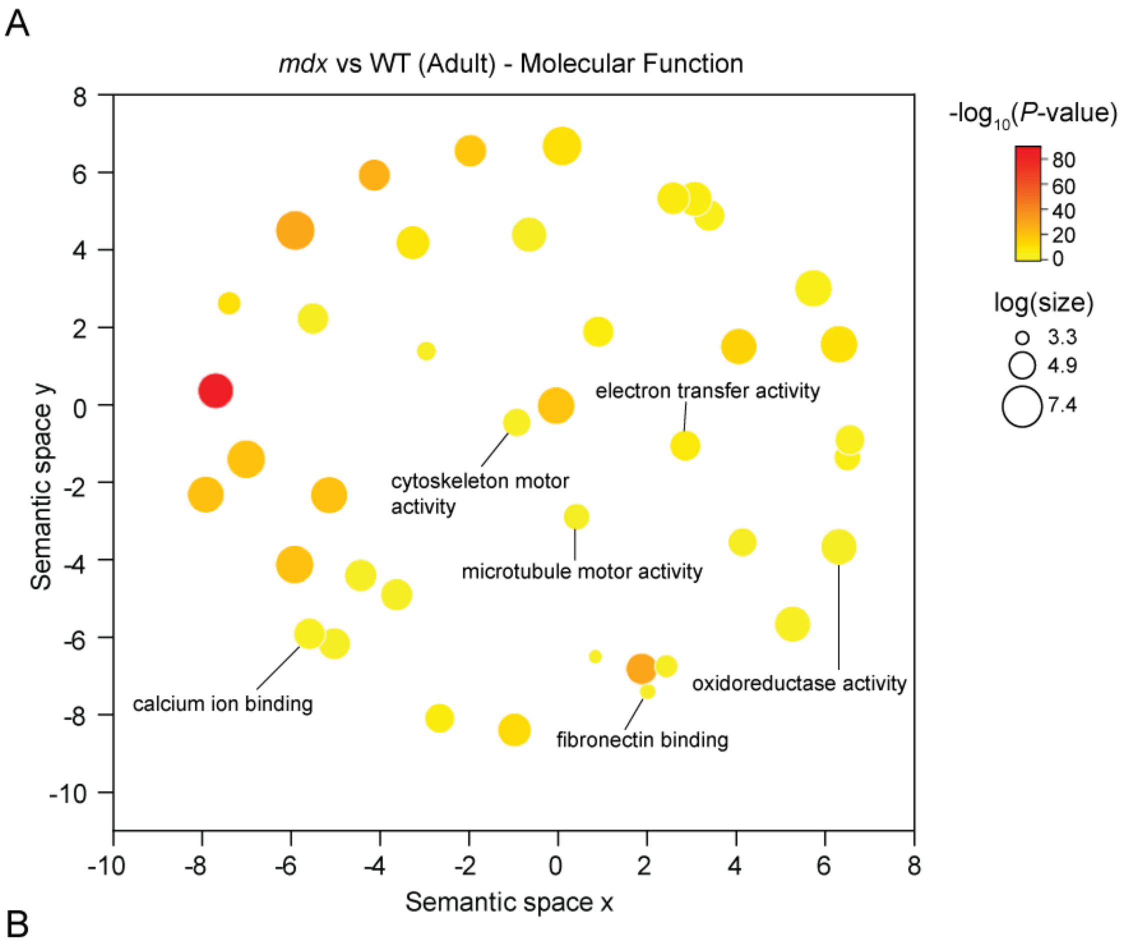
A
mdx vs WT (Adult) - Molecular Function
-log10(P-value)
80
60
40
20
0
log(size)
3.3
4.9
7.4
Semantic space y
Semantic space x
electron transfer activity
cytoskeleton motor activity
microtubule motor activity
oxidoreductase activity
calcium ion binding
fibronectin binding

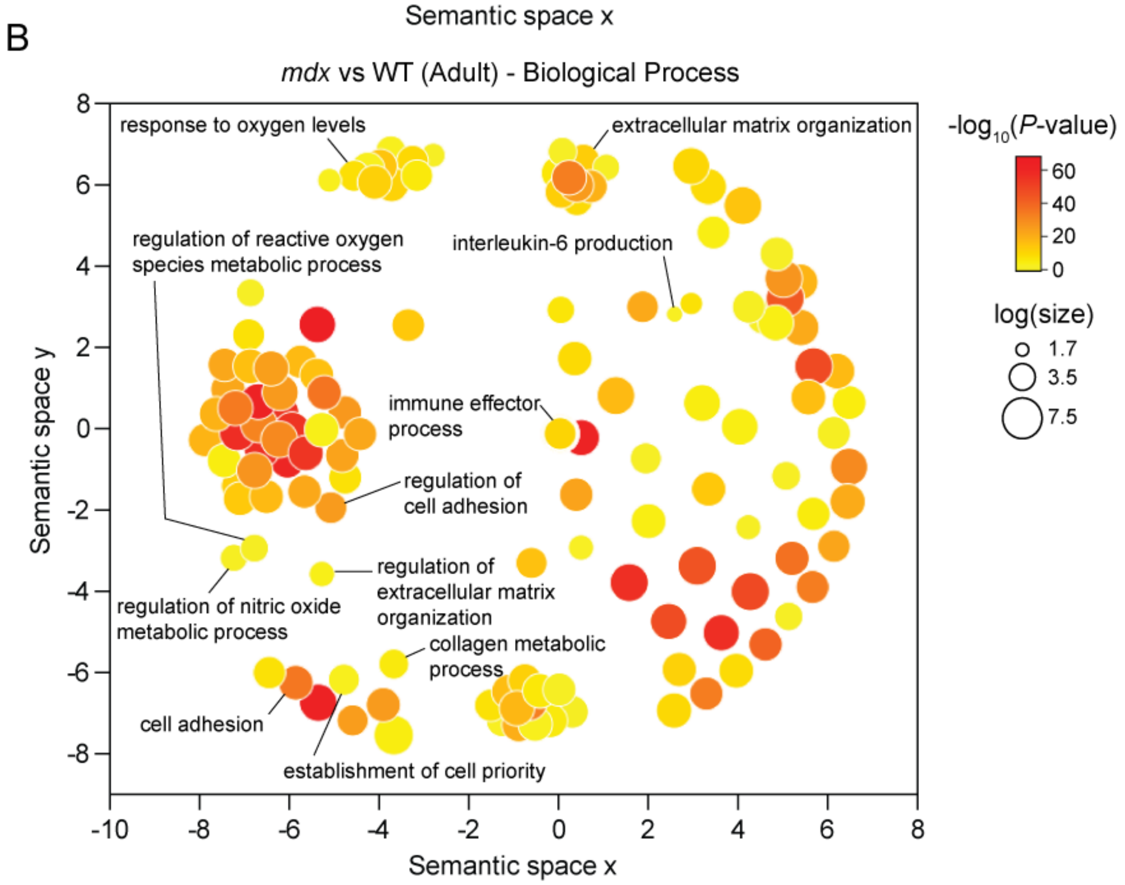
B
mdx vs WT (Adult) - Biological Process
-log10(P-value)
60
40
20
0
log(size)
1.7
3.5
7.5
Semantic space y
Semantic space x
response to oxygen levels
extracellular matrix organization
regulation of reactive oxygen species metabolic process
interleukin-6 production
immune effector process
regulation of cell adhesion
regulation of nitric oxide metabolic process
regulation of extracellular matrix organization
collagen metabolic process
cell adhesion
establishment of cell priority

**Figure S2**

**Gene ontology analysis for differentially expressed genes in Adult *mdx* vs WT.**

Differentially-expressed genes from RNA-Seq data (adjusted $P<0.01$) for the Adult *mdx* vs WT libraries were analysed for enriched gene ontology (GO) terms using g:Profiler and the statistically significant (adjusted $P<0.05$) terms visualized using REVIGO to reduce redundancy and improve interpretability for (**A**) molecular function, and (**B**) biological process. Circle size indicates the frequency of the gene ontology annotation term in the underlying database, and colour indicates the statistical significance of the enrichment. Distances between circles reflect semantic similarity among GO terms.

A

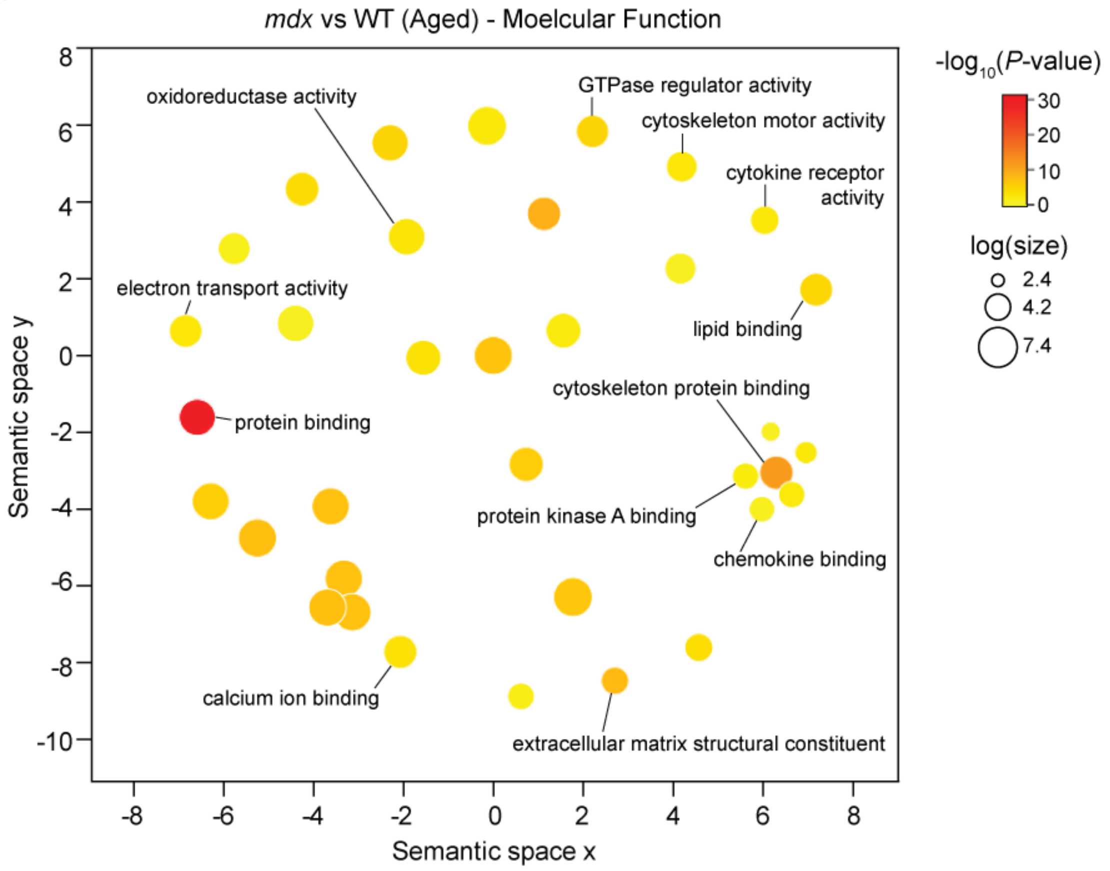
mdx vs WT (Aged) - Moelcular Function
oxidoreductase activity
GTPase regulator activity
cytoskeleton motor activity
cytokine receptor activity
electron transport activity
lipid binding
cytoskeleton protein binding
protein binding
protein kinase A binding
chemokine binding
calcium ion binding
extracellular matrix structural constituent
Semantic space y
Semantic space x
-log10(P-value)
log(size)
2.4
4.2
7.4

B

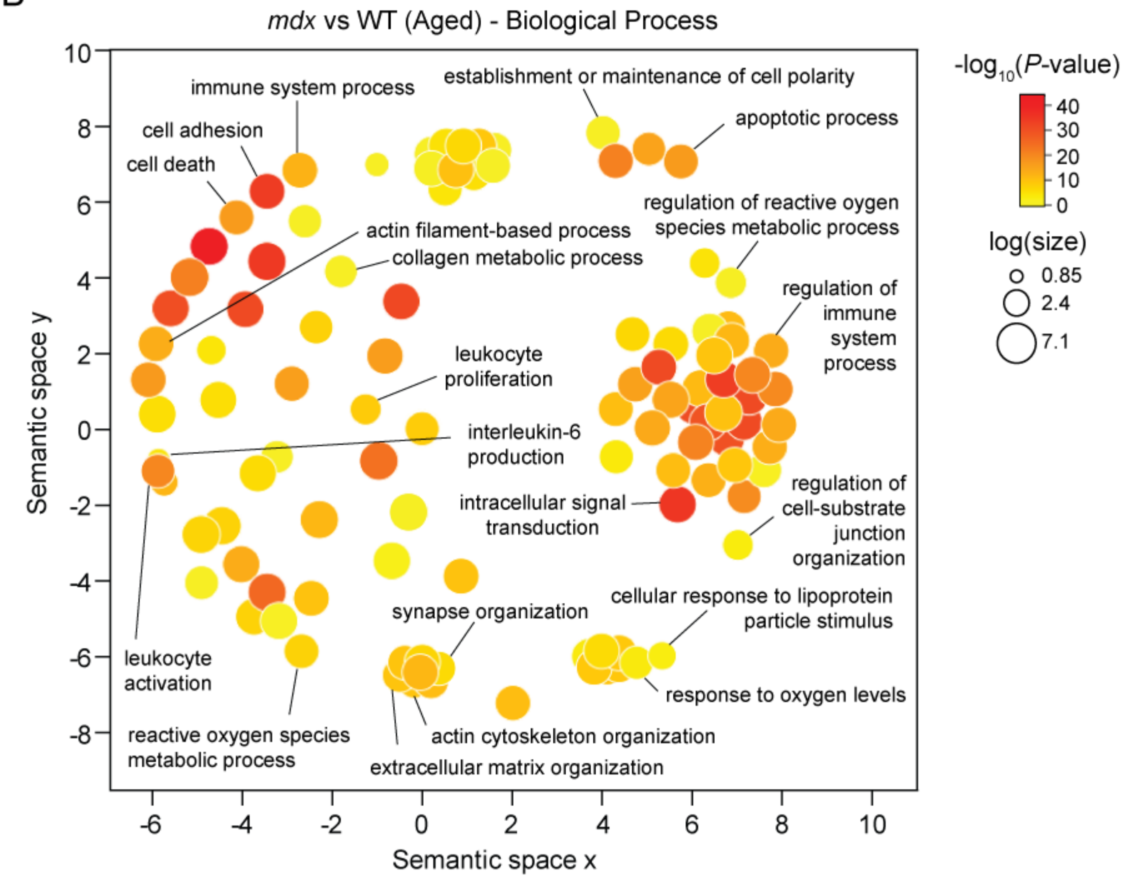
mdx vs WT (Aged) - Biological Process
immune system process
establishment or maintenance of cell polarity
apoptotic process
cell adhesion
cell death
actin filament-based process
collagen metabolic process
regulation of reactive oxygen species metabolic process
regulation of immune system process
leukocyte proliferation
interleukin-6 production
intracellular signal transduction
regulation of cell-substrate junction organization
synapse organization
cellular response to lipoprotein particle stimulus
leukocyte activation
response to oxygen levels
reactive oxygen species metabolic process
actin cytoskeleton organization
extracellular matrix organization
Semantic space y
Semantic space x
-log10(P-value)
log(size)
0.85
2.4
7.1

**Figure S3**

**Gene ontology analysis for differentially expressed genes in Aged *mdx* vs WT.**

Differentially-expressed genes from RNA-Seq data (adjusted $P<0.01$) for the Aged *mdx* vs WT libraries were analysed for enriched gene ontology (GO) terms using g:Profiler and the statistically significant (adjusted $P<0.05$) terms visualized using REVIGO to reduce redundancy and improve interpretability for (**A**) molecular function, and (**B**) biological process. Circle size indicates the frequency of the gene ontology annotation term in the underlying database, and colour indicates the statistical significance of the enrichment. Distances between circles reflect semantic similarity among GO terms.

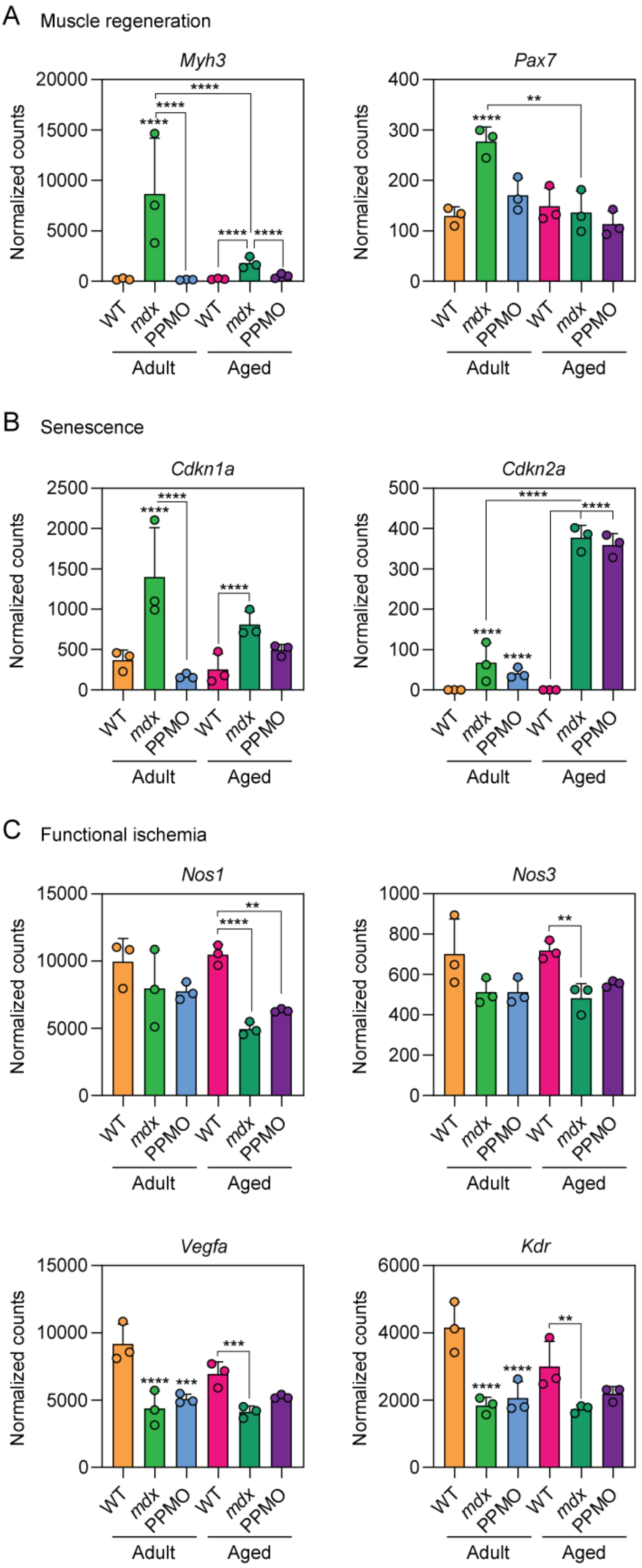
A Muscle regeneration
Myh3
Pax7
B Senescence
Cdkn1a
Cdkn2a
C Functional ischemia
Nos1
Nos3
Vegfa
Kdr
Normalized counts
WT
mdx
PPMO
Adult
Aged

**Figure S4**

**Differential expression of muscle regeneration, senescence, and functional ischemia markers in dystrophic muscle.**

RNA-Seq normalized counts data for marker genes associated with (**A**) muscle regeneration, (**B**) senescence, and (**C**) functional ischemia. Values are mean+SD. Statistical significance was assessed using DESeq2 and Benjamini-Hochberg-adjusted *P*-values indicated. Statistical comparisons are relative to the Adult WT group unless otherwise indicated, **$P<0.01$, ***$P<0.001$, ****$P<0.0001$.

Handling of reactive oxygen species

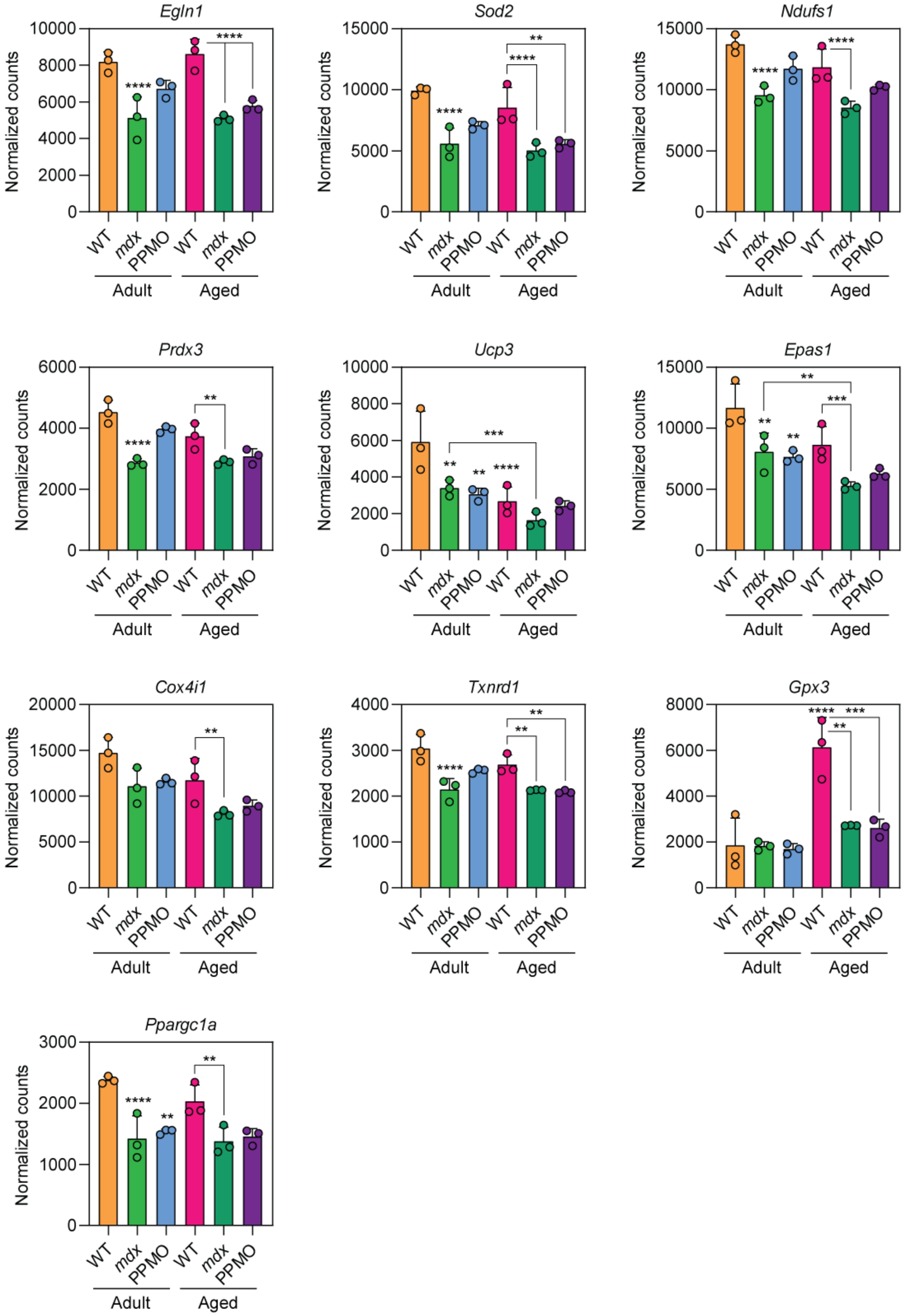

**Figure S5**

**Differential expression of reactive oxygen species handling markers in dystrophic muscle.**

RNA-Seq normalized counts data for marker genes associated with handling of reactive oxygen species. Values are mean+SD. Statistical significance was assessed using DESeq2 and Benjamini-Hochberg-adjusted *P*-values indicated. Statistical comparisons are relative to the Adult WT group unless otherwise indicated, $^{**}P<0.01$, $^{***}P<0.001$, $^{****}P<0.0001$.

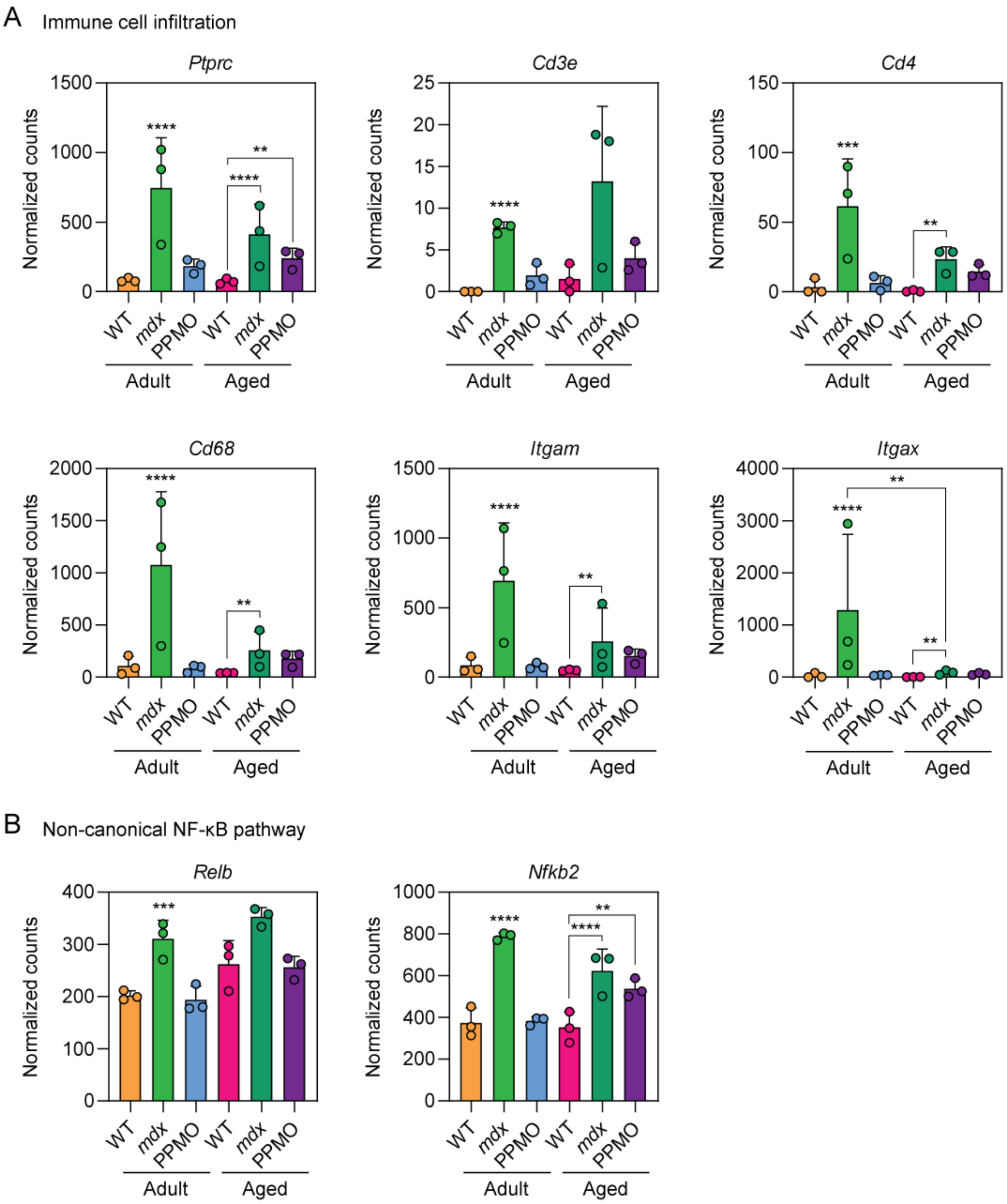
A
Immune cell infiltration
Ptprc
Cd3e
Cd4
Cd68
Itgam
Itgax
Normalized counts
0
500
1000
1500
5
10
15
20
25
50
100
150
2000
3000
4000
WT
mdx
PPMO
Adult
Aged
****
***
**
B
Non-canonical NF-κB pathway
Relb
Nfkb2
100
200
300
400
600
800

**Figure S6**

**Differential expression of immune infiltration and NF-κB pathway markers in dystrophic muscle.** RNA-Seq normalized counts data for marker genes associated with (**A**) immune cell infiltration and (**B**) the non-canonical NF-κB pathway. Values are mean+SD. Statistical significance was assessed using DESeq2 and Benjamini-Hochberg-adjusted *P*-values indicated. Statistical comparisons are relative to the Adult WT group unless otherwise indicated, ***P*<0.01, ****P*<0.001, *****P*<0.0001.

Fibrosis

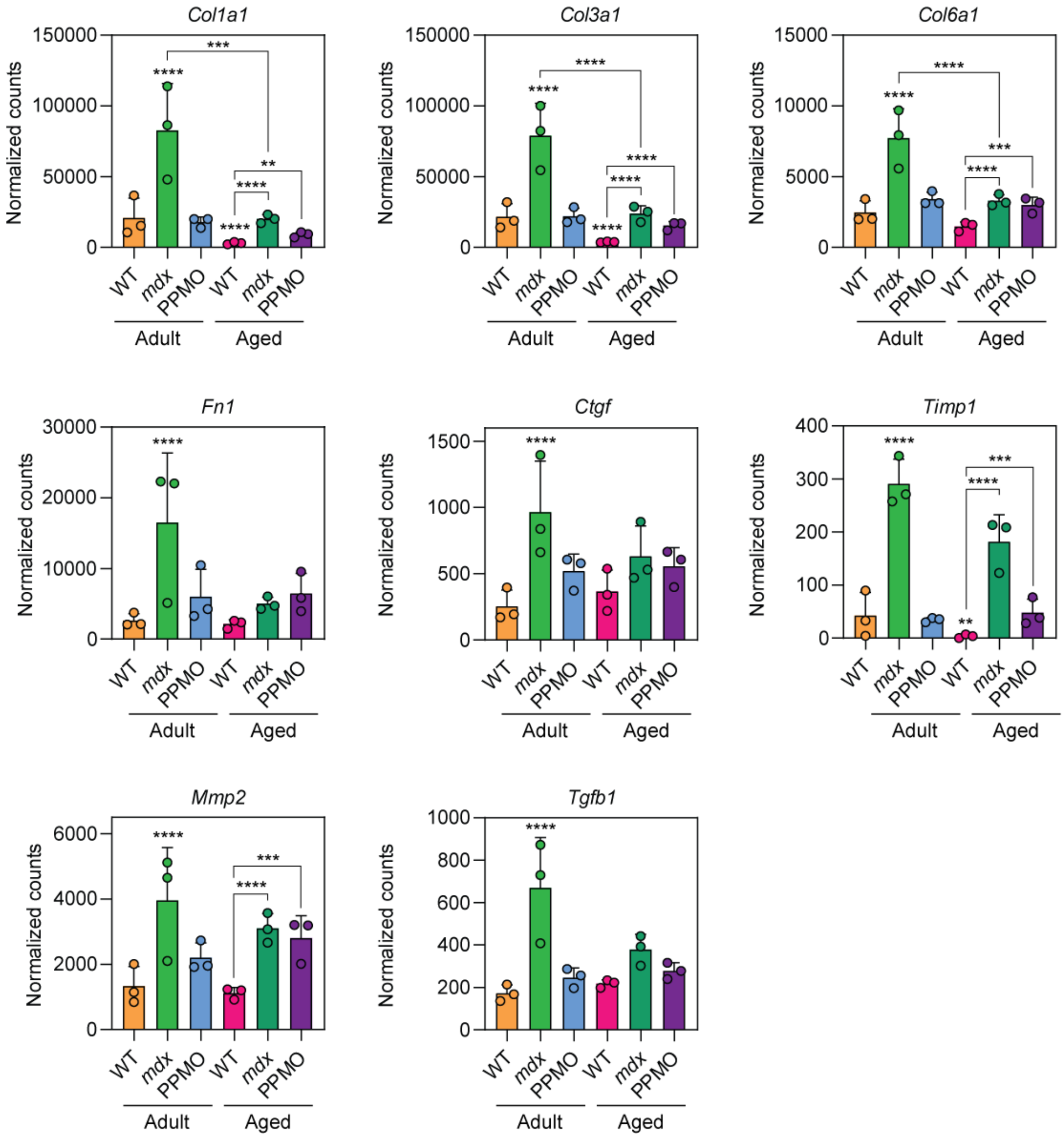

Col1a1
Col3a1
Col6a1
Fn1
Ctgf
Timp1
Mmp2
Tgfb1
Normalized counts
WT
mdx
PPMO
Adult
Aged
****
***
**

**Figure S7**

**Differential expression of fibrosis markers in dystrophic muscle.**

RNA-Seq normalized counts data for marker genes associated with fibrosis. Values are mean+SD. Statistical significance was assessed using DESeq2 and Benjamini-Hochberg-adjusted *P*-values indicated. Statistical comparisons are relative to the Adult WT group unless otherwise indicated, $^{**}P<0.01$, $^{***}P<0.001$, $^{****}P<0.0001$.

AKT/PI3K/mTOR axis

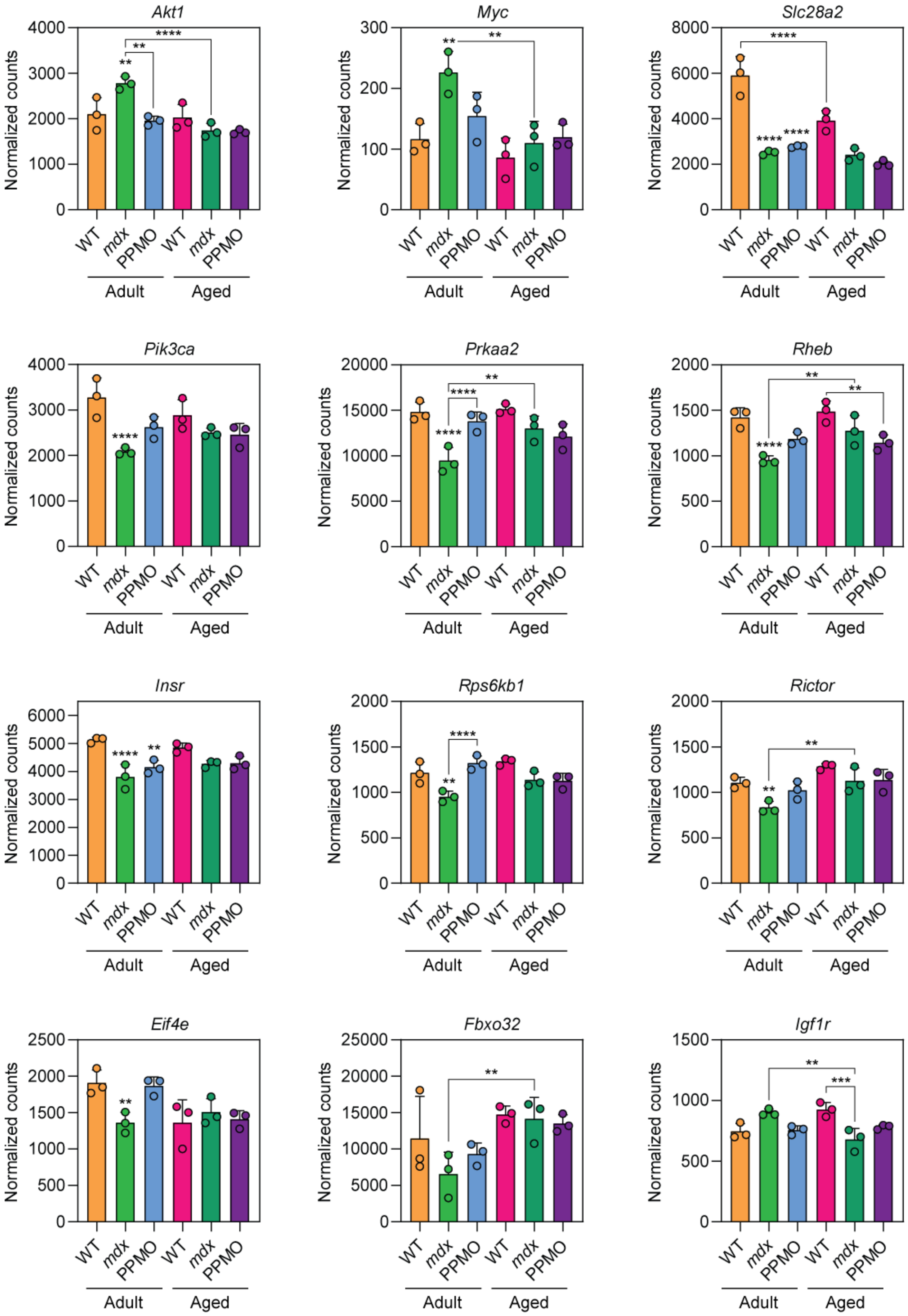

**Figure S8**

**Differential expression of AKT/PI3K/mTOR axis markers in dystrophic muscle.**

RNA-Seq normalized counts data for marker genes associated with the AKT/PI3K/mTOR axis. Values are mean+SD. Statistical significance was assessed using DESeq2 and Benjamini-Hochberg-adjusted *P*-values indicated. Statistical comparisons are relative to the Adult WT group unless otherwise indicated, $^{**}P<0.01$, $^{***}P<0.001$, $^{****}P<0.0001$.
